# Fitness at the Expanding Front: An Exploration-Exploitation Trade-off in Phenotypic Switching

**DOI:** 10.1101/2025.10.27.684987

**Authors:** Vedant Sharma, Hao Wang

## Abstract

Phenotypic switching is a key bet-hedging strategy for navigating the exploration-exploitation trade-off in fluctuating environments, yet its interplay with spatial population dynamics during range expansions remains poorly understood. We use an individual-based spatial simulation to investigate how switching strategies shape fitness (expansion speed) in heterogeneous landscapes. Maximizing expansion speed requires balancing exploration and exploitation, leading to an optimal intermediate switching rate in these spatial settings. Critically, we incorporate a phenotypic switching lag, representing the biophysical cost of adaptation. We demonstrate this lag imposes a hard fitness constraint, forcing the optimal strategy toward slower switching rates as lag duration increases and reducing maximum expansion speed. Contrasting with models focusing on irreversible mutations, we show reversible switching enhances resilience by allowing recovery from maladaptation, influencing specialist persistence boundaries. Spatial structure also generates emergent phenomena: we find clustering provides collective protection, enhancing the survival of disadvantaged phenotypes. Additionally, our simulations show that conditioning on lineage survival alone is sufficient to generate the apparent stagnancy of deleterious sectors reported experimentally, revealing this observational bias as a crucial factor when characterizing selection effects at expanding fronts. This work integrates bet-hedging theory with adaptation costs and spatial dynamics, offering a quantitative framework for fitness at expanding fronts.

## Introduction

How populations adapt to fluctuating environments is a central question in evolutionary biology. Environmental conditions, from nutrient availability to antibiotic presence, are rarely stable, demanding strategies that ensure long-term survival. Phenotypic switching, where genetically identical individuals stochastically adopt distinct phenotypes, is a key bet-hedging mechanism allowing populations to spread risk across generations (16). Driven by noise in gene regulation, this non-genetic diversity ensures some individuals are prepared for unforeseen challenges, as seen in bacterial persistence (3) and anticipatory metabolic shifts in yeast (20).

Such strategies inherently involve an exploration-exploitation trade-off. Minimizing switching (exploitation) maximizes growth under current conditions but increases vulnerability to change. Conversely, frequent switching (exploration) prepares for future shifts but incurs a constant fitness cost from maladapted individuals (18). The optimal balance, often characterized by an intermediate switching rate, depends on the environment’s fluctuation frequency, typically studied in well-mixed systems (1).

However, many biological populations, from microbial colonies to invading species, are spatially structured and expand their range (7, 16). This spatial context fundamentally alters population dynamics compared to well-mixed assumptions. At expanding fronts, genetic drift is amplified, allowing even deleterious traits to “surf” to high frequencies (7), while collective cellular interactions can buffer individuals from selection (2, 11). Furthermore, environmental heterogeneity can make location more critical than intrinsic fitness (6). It thus remains unclear how temporal bet-hedging strategies via phenotypic switching perform when confronted with the altered rules of selection and drift at an expanding front in a heterogeneous landscape.

Here, we address this gap using a stochastic, individual-based spatial simulation (specifically, a Gillespie-based Eden model on a hexagonal lattice) to investigate the strategic landscape of phenotypic switching during range expansion. Our investigation first establishes the fitness landscape (measured by front speed), confirming an optimal intermediate switching rate balances exploration and exploitation in this spatial context. We then analyze the strategic importance of switching reversibility, contrasting its effects on population resilience and persistence boundaries with irreversible strategies often studied in mutation models (12). A key focus is incorporating the biophysical cost of adaptation via a phenotypic switching lag (4, 9), revealing its significant constraint on fitness and its role in shifting optimal strategies. Finally, we examine emergent spatial phenomena, demonstrating collective protection through clustering and showing how conditioning on lineage survival provides a mechanistic explanation for counter-intuitive observations like the apparent stagnancy of deleterious sectors (2, 14).

## Results

### An Optimal Intermediate Switching Rate Maximizes Front Speed

We quantified long-term population fitness by the mean speed of the expanding front (*v*_*f*_), a suitable measure in our spatially and temporally fluctuating environment (17). Simulating populations with symmetric switching between phenotypes across a range of total switching rates (*k*_*total*_), we found that fitness varies non-monotonically with the switching rate (Fig. 1A). Front speed is maximized at an optimal intermediate rate, *k*_*opt*_, decreasing substantially at both very low (*k*_*total*_ ≪ *k*_*opt*_) and very high (*k*_*total*_ ≫ *k*_*opt*_) switching rates.

**Fig. 1.**
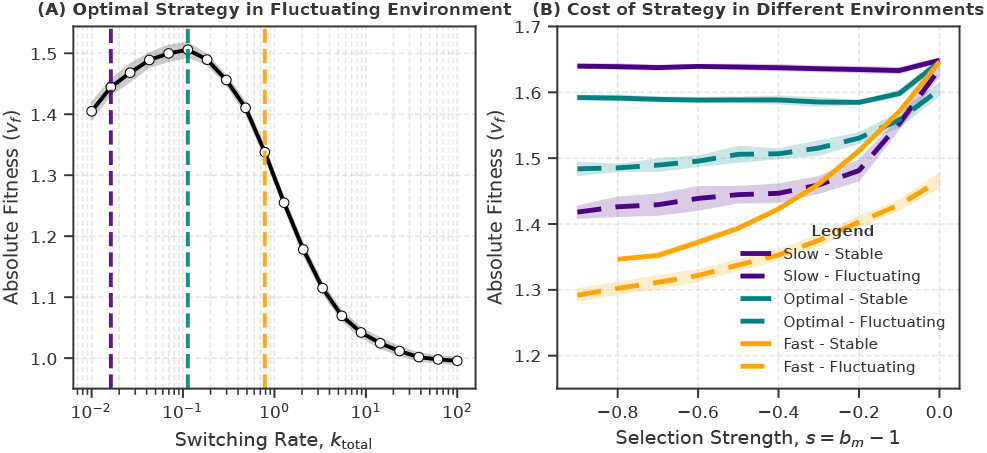
The Bet-Hedging Trade-Off. (A) Absolute fitness (front speed) as a function of the total switching rate, *k*_total_, in a fluctuating environment. Fitness is maximized at an optimal intermediate rate. (B) Relative fitness of slow, optimal, and fast switching strategies under different selection strengths in stable (solid) and fluctuating (dashed) environments.

To further characterize these strategies, we compared the performance of representative slow, optimal, and fast switching rates under varying selection strengths in both stable and fluctuating environments (Fig. 1B). In a stable environment where the specialist phenotype is always disadvantageous (solid lines), fitness generally decreased with increasing switching rate, indicating a cost to switching. In contrast, within the fluctuating environment (dashed lines), the optimal switching strategy maintained the highest fitness across strong negative selection pressures, while the slow-switching strategy performed poorly under such conditions.

### Switching Reversibility Enhances Fitness and Alters Persistence Boundaries

Beyond the total switching rate, the directionality of switching represents a distinct strategic dimension. We investigated this using a switching bias parameter, *ϕ*, which partitions *k*_*total*_ into forward (*k*_*G*→*S*_) and reverse (*k*_*S*→*G*_) rates, The parameter *ϕ* ranges from *ϕ* = −1.0 (irreversible G→S switching) to *ϕ <* 1.0 (allowing reversible S→G switching). The case *ϕ* = 1.0 (no S→G switching) was excluded as it leads to rapid extinction in hostile environments.

First, we examined a scenario analogous to diauxic growth (20), where the specialist phenotype is highly fit in its preferred environment (*b*_*m*_ = 1.0) but less fit elsewhere. In simulations within a periodically fluctuating environment, reversible strategies (*ϕ >* −1.0) consistently achieved higher long-term fitness (front speed) compared to the irreversible strategy (*ϕ* = −1.0), particularly under stronger selection against the specialist in the non-preferred patch (Fig. 2A-C). Plotting the fitness achieved by the optimal strategy (maximized over *k*_*total*_ and *ϕ* for each *b*_*m*_) against the irreversible strategy highlights this “Advantage of Reversibility”, which increases as the specialist’s fitness cost grows (Fig. 2D).

**Fig. 2.**
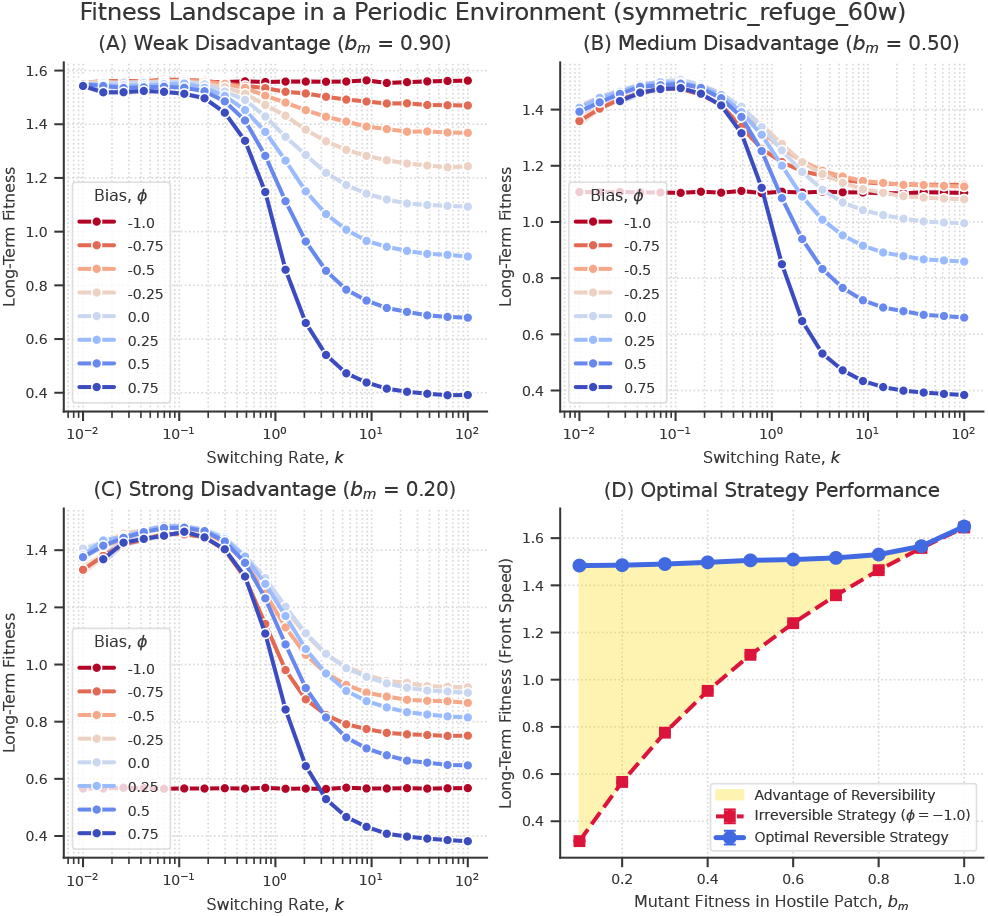
Fitness Landscape in a Periodic Environment. (A-C) Long-term fitness as a function of switching rate for various switching biases (*ϕ*) and specialist fitness in hostile patches (*b*_*m*_). Reversible strategies (*ϕ >* −1.0, blue-toned lines) consistently outperform the irreversible strategy (*ϕ* = −1.0, red line). (D) Performance of the optimal strategy for each condition. The yellow shaded area highlights the “Advantage of Reversibility,” the fitness gained by allowing back-switching from the specialist state.

To further investigate the role of reversibility, we analyzed population stability in a homogeneous, hostile environment where the specialist phenotype is always disadvantageous (*b*_*m*_ *<* 1.0). We mapped the final steady-state specialist fraction ⟨*ρ*_*M*_ ⟩ as a function of selection strength (*s* = *b*_*m*_ − 1.0) and switching rate (*k*_*total*_) for different bias values (Fig. 3A-C). These phase diagrams show a transition between an active phase (mixed population) and an absorbing phase (specialists eliminated or fixed, depending on interpretation and initial conditions relative to *s*). The location of the boundary between these phases depends strongly on the switching bias *ϕ*. Strategies with greater reversibility (higher *ϕ*) allow the mixed population to persist under stronger negative selection (more negative *s*) or at lower switching rates for a given *s*, shifting the phase boundary accordingly (Fig. 3D).

**Fig. 3.**
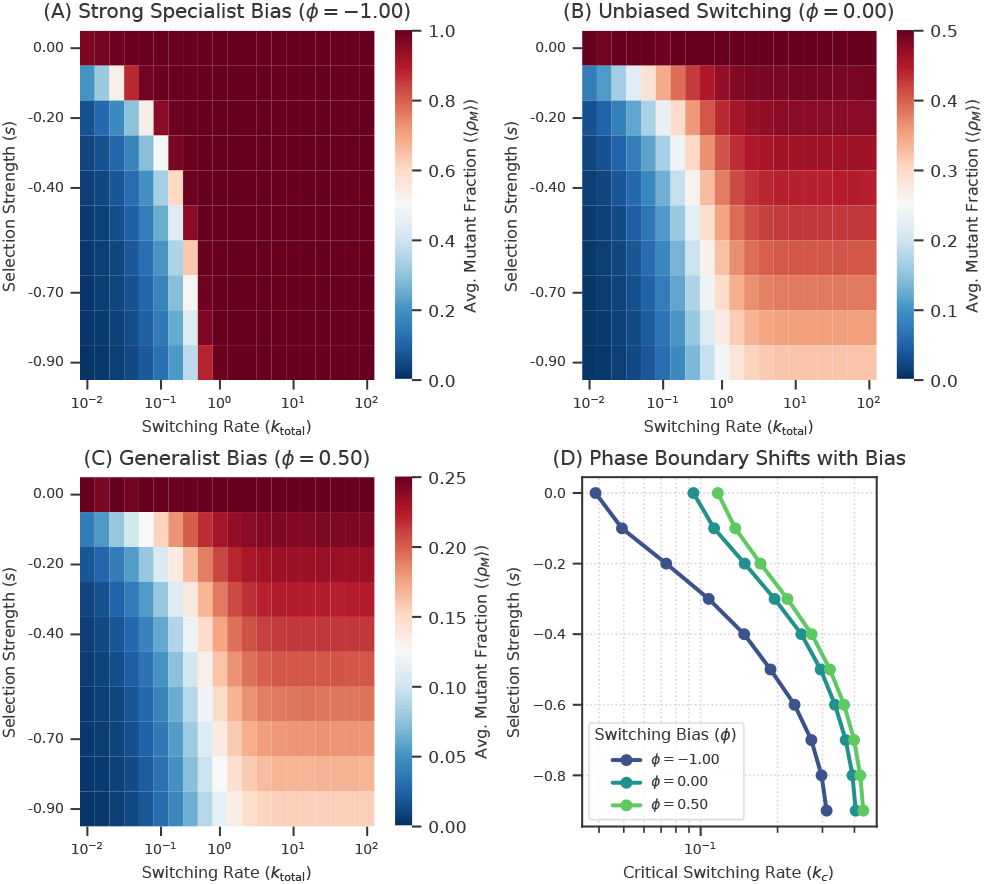
Switching bias regulates the conditions for specialist persistence. (A-C) Phase diagrams showing the final specialist fraction as a function of selection strength (*s*) and switching rate (*k*_total_) for three different switching biases (*ϕ*). The transition from an active (mixed generalist/specialist, purple) to an absorbing (all-specialist, yellow) state depends critically on the bias. (D) The phase boundary shifts to higher critical switching rates as the strategy becomes more reversible.

The steady-state specialist fraction in the active phase can be approximated by a mean-field model (refer Materials and Methods), which accurately captures the simulation results when selection is weak, though it deviates under strong selection and does not describe transient dynamics (Fig. S4). We also confirmed the advantage of reversibility in a “resilience” trade-off scenario, where the specialist incurs an inherent fitness cost (*b*_*m*_ *<* 1.0) in all environments (Fig. S2). While quantitatively smaller, the qualitative result holds: reversible strategies outperform irreversible ones (Fig. S2D).

### The Biological Cost of Adaptation Constrains Optimal Strategy

Phenotypic switching involves complex molecular reprogramming that takes time (4, 9, 10, 13). To investigate the impact of this inherent delay, we introduced a switching lag duration (*τ*) into the model, representing a period where switching cells become temporarily non-reproductive.

Incorporating this lag significantly reshapes the fitness landscape (Fig. 4A). For any fixed switching rate (*k*_*total*_), increasing the lag duration *τ* substantially reduces the population fitness (front speed). Conversely, for a fixed lag duration, fitness still varies non-monotonically with *k*_*total*_, exhibiting an optimal intermediate rate *k*_*opt*_. Notably, the location of this optimum shifts systematically: as the lag duration *τ* increases, the optimal switching rate *k*_*opt*_ decreases (indicated by the diagonal structure of the high-fitness ridge in Fig. 4A). Furthermore, the maximum achievable fitness (the fitness obtained at *k*_*opt*_ for a given *τ*) is sharply constrained by the lag duration (Fig. 4B). Longer lag durations lead to a dramatic reduction in the highest possible front speed, with the effect being most pronounced under stronger selection against the specialist phenotype (lower *b*_*m*_).

**Fig. 4.**
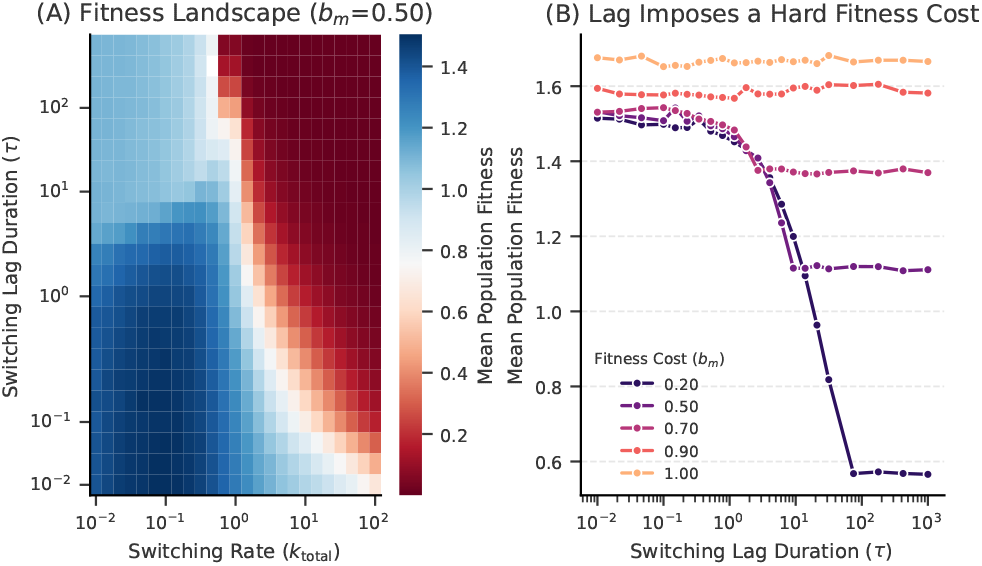
The intrinsic biological cost of adaptation reshapes the fitness landscape and optimal strategy. (A) Fitness landscape showing the mean front speed as a function of the switching lag duration and the total switching rate (*k*_total_) for a specialist fitness of *b*_*m*_ = 0.50. The ridge of high fitness (warm colors) systematically shifts to lower switching rates as the lag duration increases, indicating a shift in the optimal strategy. (B) Maximum achievable fitness (the fitness at the optimal switching rate for each condition) as a function of the switching lag duration. The lag imposes a hard fitness cost that is most severe for specialists under strong selection (low *b*_*m*_).

### Spatial Organization Creates Emergent Collective Protection

The spatial arrangement of individuals, a factor absent in well-mixed models, can significantly influence evolutionary outcomes during range expansions. We investigated this by simulating the expansion of populations starting with a fixed number of disadvantaged specialist cells (*b*_*m*_ *<* 1.0) arranged in patterns of varying spatial clustering, generated using a correlated Gaussian random field method (Fig. S1). We found that the initial spatial arrangement strongly affects the persistence of the specialist lineage. Specifically, increased clustering enhances survival. The mean peak invasion depth reached by the specialist lineage before extinction increases monotonically with the mean initial cluster size (Fig. 5A). This effect is particularly pronounced under strong negative selection; normalizing the invasion depth by that of the most fragmented state reveals that the relative benefit of clustering increases dramatically as selection against the specialist becomes stronger (Fig. 5B). For instance, under strong selection (*b*_*m*_ = 0.7), a highly clustered population persists significantly longer and invades deeper than a fragmented population with the same initial number of specialists (Fig. S3).

**Fig. 5.**
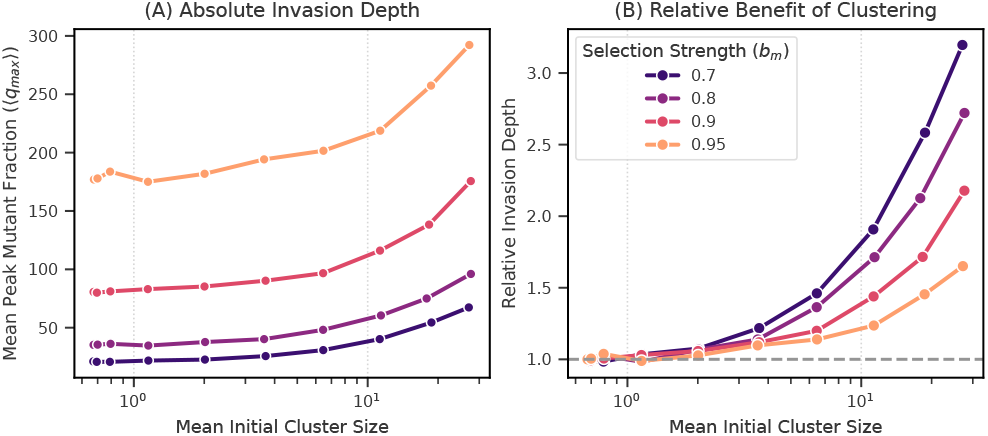
Spatial clustering enhances the persistence of disadvantaged specialist lineages. (A) Absolute invasion depth of a mutant lineage before extinction as a function of its mean initial cluster size, for different selection strengths (*b*_*m*_). (B) Relative benefit of clustering, normalized to the most fragmented state (size = 1). Clustering provides a dramatic survival advantage that increases with selection strength.

### Conditioning on Survival Explains Apparent Stagnancy of Deleterious Patches

The stochastic nature of extinction at expanding fronts can lead to observational biases when tracking lineage dynamics. Specifically, the apparent stagnancy or slow shrinkage of deleterious mutant sectors observed experimentally (2) can emerge from conditioning observations on lineage survival. We investigated this phenomenon using our simulation framework in both linear and radial expansion geometries (Fig. 6).

**Fig. 6.**
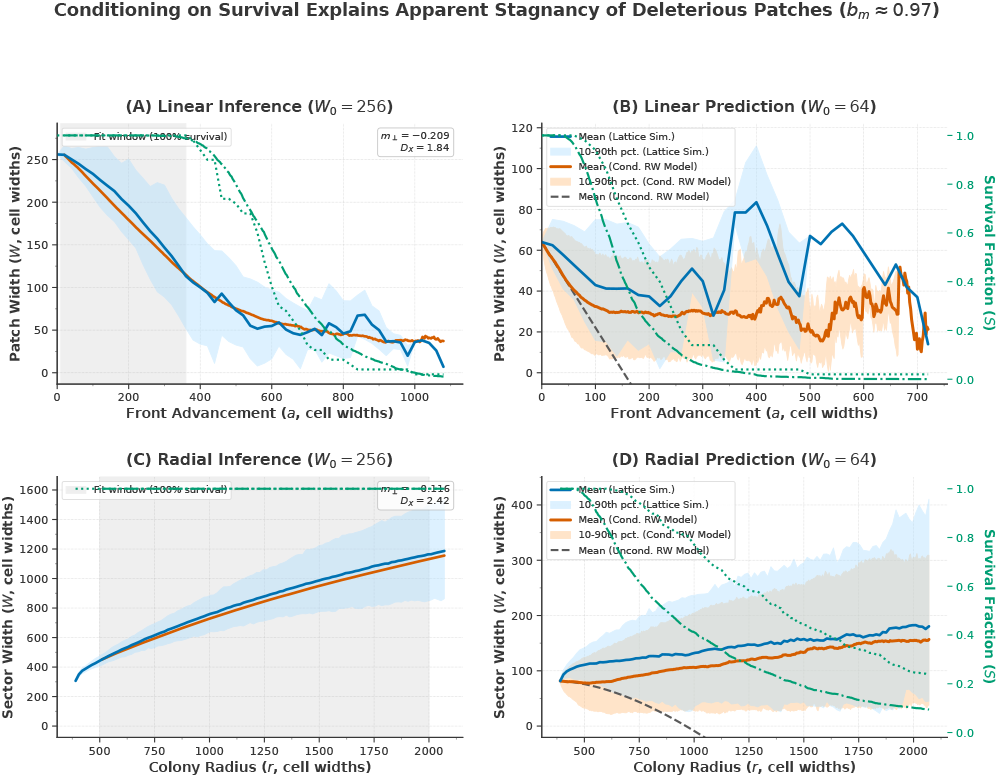
Conditioning on survival explains the apparent stagnancy of deleterious patches in both linear and radial geometries. We compare data from our full agent-based “Lattice Sim” (blue/light blue) against a simplified “Conditioned Random Walk (RW) Model” (orange/light orange), using parameters inferred from one dataset (*W*_0_ = 256) to predict another (*W*_0_ = 32). The mutant fitness *b*_*m*_ is held constant at ≈ 0.97 (*s* ≈ −0.03). Details of the RW model and parameter inference are in the Methods section. **(A) Linear Inference (***W*_0_ = 256): Parameters *m*_⊥_ and *D*_*X*_ are inferred from the lattice simulation within the high-survival window. The conditioned RW model (orange) closely follows the lattice data (blue). **(B) Linear Prediction (***W*_0_ = 32): Using parameters from (A), the conditioned RW model (orange line: mean; light orange shading: 10-90th percentile) accurately predicts the mean (blue line) and distribution (light blue shading) of the lattice simulation. The apparent stagnancy contrasts sharply with the prediction of the unconditioned RW model (dashed black line). Survival fractions (*S*) for lattice (dotted green) and con-ditioned RW (dash-dot green) are shown on the right axis. **(C) Radial Inference (***W*_0_ = 256): The same inference procedure applied to radial expansion yields different effective *m*_⊥_ and *D*_*X*_ values. **(D) Radial Prediction (***W*_0_ = 32): Parameters from (C) accurately predict the dynamics for the smaller initial width in the radial case. In both (B) and (D), the conditioned RW model correctly captures the apparent stagnancy caused by observing only surviving lineages

First, we inferred the effective parameters for a biased random walk (RW) model—drift *m*_⊥_ and diffusion *D*_*X*_ —from agent-based simulations initiated with a large patch width (*W*_0_ = 256 cells), where initial survival probability is high, Fig. 6 A, C, Fig. S7). These parameters quantify the average shrinkage rate due to selection (*m*_⊥_ *<* 0) and the stochastic wandering of the patch boundaries (*D*_*X*_) within the highsurvival regime.

Next, using these inferred parameters without further adjustment, we compared the predictions of the simplified RW model (simulated numerically, incorporating the absorbing boundary at *W* = 0) against independent agent-based lattice simulations initiated with a smaller, more drift-sensitive initial width (*W*_0_ = 64 cells). We found excellent agreement between the full lattice simulation results (mean and distribution of patch widths for surviving lineages) and the predictions of the **conditioned** RW model across both linear (Fig. 6B) and radial (Fig. 6D) geometries. Both the lattice simulation and the conditioned RW model exhibit the apparent stagnancy or slow shrinkage reported experimentally. Crucially, these dynamics diverge significantly from the prediction of an **unconditioned** RW model (dashed black lines), which represents the theoretical average over all possible trajectories including those going extinct and shows rapid shrinkage driven by negative selection (*m*_⊥_). Furthermore, the survival fraction over time (or radius) in the lattice simulations closely matches that predicted by the conditioned RW model (Fig. 6 B, D, right axes).

## Discussion

Phenotypic switching is a well-established bet-hedging strategy for populations in temporally fluctuating environments (1, 16). However, most theory assumes well-mixed populations (19). Our work investigates this strategy in the distinct context of a spatially expanding front, where dynamics like genetic drift and local competition are fundamentally different (6, 7, 11). We find that while an optimal switching rate exists (Fig. 1), the strategy’s success is critically determined by spatial factors and biophysical costs, including the ability to reverse a switch (Fig. 2, 3), the time cost of switching (Fig. 4), and emergent spatial interactions (Fig. 5, 6).

Our spatial simulations confirm the expectation from well-mixed models (1) that fitness is maximized at an optimal, intermediate switching rate *k*_*opt*_ (Fig. 1A). This optimum arises from the fundamental exploration-exploitation trade-off, manifesting uniquely at the expanding front. Strategies with very slow switching are too ‘exploitative’; while effective within a favorable environmental patch, they fail to generate sufficient phenotypic diversity to adapt quickly when the environment changes. This leads to population stalling at patch boundaries and reduces the overall average front speed. Conversely, strategies with very fast switching are too ‘exploratory’; they constantly produce diverse phenotypes but incur a substantial fitness cost because a large fraction of cells are maladapted to the current local environment at any given time (Fig. 1B). The optimal strategy *k*_*opt*_ represents the best compromise in this spatial context.

A central finding is the strategic importance of switching reversibility. An irreversible commitment (*ϕ* = −1.0) proves fragile, yielding lower fitness in fluctuating environments (Fig. 2). This fragility arises because irreversible switches create a persistent fitness drag by generating specialists that cannot revert when the environment becomes unfavorable (12). In contrast, reversible switching (*k*_*S*→*G*_ *>* 0) provides population resilience by acting as an escape route from maladaptation. This back-switching acts as an intrinsic restoring force that counteracts negative selection, allowing the population to persist under much harsher conditions (Fig. 3A-C) and consequently shifting the phase boundary for specialist persistence (Fig. 3D).

Our analysis highlights the critical impact of the phenotypic switching lag (*τ*), an intrinsic cost of adaptation often overlooked in theoretical models (1, 19) but grounded in cellular biophysics (4, 13). Incorporating this delay reshapes the fitness landscape (Fig. 4A). The lag imposes a direct fitness cost because switching individuals are temporarily removed from the reproducing population at the front. This opportunity cost sharply constrains the maximum achievable front speed (Fig. 4B). Consequently, the optimal strategy adapts: as switching becomes more costly in terms of lost reproductive time, slower, more conservative switching rates (*k*_*opt*_ decreases as *τ* increases) are favored. This strategic shift reflects a greater emphasis on exploiting the current state rather than risking the prolonged, non-reproductive transition.

Spatial structure also generates emergent phenomena. First, we find that clustering provides **collective protection** (Fig. 5). This arises because selection acts primarily at the interface between phenotypes; clustering minimizes the boundary-to-area ratio, shielding interior cells and enhancing persistence (2, 11). Second, our model mechanistically explains the **apparent stagnancy** of deleterious sectors seen in experiments (2, 14). We show this is an observational bias (Fig. 6). The theoretical **unconditioned** mean (including extinct lineages) shrinks as expected, but the **conditioned** mean (survivors only) appears stagnant. This effect arises purely from conditioning on survival near an absorbing extinction boundary, highlighting a crucial factor in interpreting selection from experimental data (15).

While our model provides a general framework, its simplifying assumptions suggest future directions. We modeled discrete environmental patches with fixed durations, whereas real-world habitats often feature continuous gradients (6) or fluctuations drawn from a distribution. Similarly, we implemented a fixed switching lag duration *τ*, though biological lag times often exhibit variability (4). Investigating how distributions of environmental durations and lag times interact, and whether lag distributions might evolve to match environmental statistics, presents an interesting avenue. Our model also does not explicitly couple front roughness to composition, which could be explored with field-theoretic approaches (8). Furthermore, allowing the switching parameters (*k*_*total*_, *ϕ*) themselves to evolve via mutation could reveal how populations carry information about past environments and adapt their strategies over longer timescales. Understanding how the transient population composition carries information between environmental states before reaching equilibrium also warrants further study. Despite these simplifications, our core predictions regarding the impact of rate, bias, lag, and spatial structure are directly testable using synthetic gene circuits with tunable switching in microfluidic devices (10) or by extending the model to 3D to capture biofilm or tumor geometries.

In conclusion, this work provides a unifying framework that connects single-cell strategy (defined by switching rate, bias, and lag) with population-level outcomes like fitness and stability in a spatially explicit context. By showing how these parameters interact, our work offers a new, quantitative lens for understanding the emergence of resilient and adaptive behaviors in the complex, fluctuating worlds that most organisms inhabit.

## Materials and Methods

### The Two-Phenotype Spatial Switching Model

We model a spatially expanding population using a continuous-time, event-driven simulation based on the Eden model, implemented on a two-dimensional hexagonal lattice to minimize lattice artifacts (12). Each site on the lattice can be in one of four states: Empty, Generalist (wild-type), Specialist (alternative phenotype), or Transient (a temporary, non-reproductive state entered during phenotypic switching). The population is initialized as a one-dimensional front of occupied cells of width *L* with periodic boundary conditions in the transverse direction, expanding into unoccupied territory. The system’s dynamics are advanced using a Gillespie algorithm, which stochastically selects the next event based on the rates of all possible events in the system (5). To efficiently manage the large number of possible events at the expanding front, we employ a Summed-Rate Tree data structure. This allows for logarithmic-time updates and event selection, making large-scale simulations computationally feasible.

### Event Types and Rates

Two primary types of stochastic events can occur at the active front (i.e., occupied sites adjacent to empty sites):

#### 1. Growth (Reproduction)

An occupied cell at site *h* can reproduce into an adjacent empty site *h*^'^. The rate of this event depends on the cell’s phenotype and the local environmental conditions (see Fig. S5 for calibration). For a Generalist cell, the growth rate is *b*_*wt*_; for a Specialist, it is *b*_*m*_. The values of *b*_*wt*_ and *b*_*m*_ are defined per environmental patch. In this work, the Generalist growth rate is normalized to *b*_*wt*_ = 1.0 in all environments, while the Specialist growth rate *b*_*m*_ is varied to model the effects of selection.

#### 2. Phenotypic Switching

An occupied cell at the front can stochastically switch its phenotype. The dynamics of this process are governed by two key parameters: the total switching rate, *k*_total_, and the switching bias, *ϕ*. These parameters determine the individual rates for forward (Generalist → Specialist) and reverse (Specialist → Generalist) switching according to the following relations:

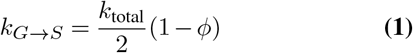

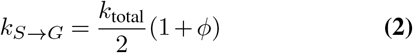

The bias parameter *ϕ* ranges from -1 to +1. A bias of *ϕ* = −1.0 corresponds to a purely irreversible “specialist” strategy (*k*_*S*→*G*_ = 0), where Generalists can become Specialists but cannot revert. A bias of *ϕ* = 0.0 represents unbiased, symmetric switching.

### Implementation of Switching Lag

To model the intrinsic biological cost of adaptation, we introduce a *switching lag duration* (*τ*), a period of phenotypic inertia where a cell becomes temporarily non-reproductive. When a switching event is selected by the Gillespie algorithm, the cell does not instantly change its phenotype. Instead, it enters a “Transient” state for a deterministic duration, *τ*. During this lag period, the cell is inert; it cannot grow or initiate further switches. This is implemented using a priority queue (min-heap) that stores scheduled “completion” events. The main simulation loop advances by choosing the event with the minimum time-to-occurrence, whether it is the next stochastic event from the Gillespie algorithm or the next deterministic completion event from the priority queue. When a completion event is executed, the cell exits the Transient state, adopts its final phenotype, and its potential for growth and switching is re-evaluated.

### Generation of Spatially Correlated Initial Conditions

To investigate the role of spatial organization, we developed a method to generate initial population fronts with a fixed number of specialist cells but with varying degrees of spatial clustering (Fig. S1). This is achieved by thresholding a one-dimensional Correlated Gaussian Random Field (GRF). The process is as follows:

1. A 1D continuous field of “suitability” values is generated along the population front of width *L*. The spatial correlation of this field is controlled by a parameter, *λ*_*c*_ (correlation length).
2. A rank-based threshold is applied. We identify the *N*_*S*_ sites with the highest field values, where *N*_*S*_ is the desired number of initial specialist cells.
3. These top *N*_*S*_ sites are designated as Specialist cells, and the remaining *L* − *N*_*S*_ sites are designated as Generalist cells. A small correlation length (*λ*_*c*_ → 0) produces a highly fragmented, nearly random pattern, while a large correlation length (*λ*_*c*_ ≫ *L*) produces a single, cohesive cluster.

### Calculation of Fitness in Fluctuating Environments

The population’s long-term fitness, Λ, is defined as the steady-state speed of the expanding front. Theoretically, this corresponds to the long-term growth rate of the population in a fluctuating environment. Because the instantaneous front speed fluctuates as the population crosses different environmental patches, we measure the cycle-averaged front speed (the average speed over one full environmental period). The simulation is run until this cycle-averaged speed converges to a steady state, defined as the point where its relative standard deviation (over a moving window of *N* cycles) falls below a 1% threshold. This stable, converged front speed *v*_*f*_ is reported as the fitness Λ.

### Conditioned Random Walk (RW) Model

To model the dynamics of a single patch width (Fig. 6), we use a simplified drift-diffusion model (a biased random walk) with an absorbing boundary at *W* = 0, following (2). The width of the patch, *W*, evolves in time (or equivalently, with front advancement *a*) according to the stochastic differential equation:

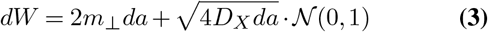

where *m*_⊥_ is the effective drift (selection) parameter, *D*_*X*_ is the effective diffusion (boundary wandering) parameter, and 𝒩 (0, 1) is a standard normal random variable.

The parameters *m*_⊥_ and *D*_*X*_ are inferred from the agent-based simulations, not assumed *a priori*. We run simulations with a large initial width *W*_0_ (e.g., *W*_0_ = 256) where extinction is rare at early times. In this high-survival window, the unconditioned mean and variance are well-approximated by:

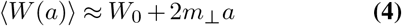

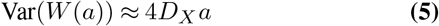

We perform linear fits on the mean width ⟨*W* (*a*)⟩ vs. *a* to find *m*_⊥_ (from the slope) and on the variance Var(*W* (*a*)) vs. *a* to find *D*_*X*_ (from the slope) (see Fig. S7 for a visualization of this inference procedure). These empirically-inferred parameters are then used in a numerical simulation of the RW model (Eq. 3) to generate the predictions (conditioned mean, percentiles, and survival fraction) shown in Fig. 6 B, D. For the radial case (Fig. 6 C, D), a change of variables Φ = *W/r* is used, leading to a similar SDE with radius-dependent coefficients, as detailed in (2) (see Fig. S6 for an illustration of radial sector dynamics).

### Mean-Field Model for Steady-State Fraction

To model the steady-state specialist fraction *ρ*_*M*_ in a homogeneous environment (Fig. 3 and Fig. S4), we use a mean-field rate equation. Assuming the fractions of generalists (*ρ*_*G*_) and specialists (*ρ*_*M*_) sum to 1, the rate of change of *ρ*_*M*_ is given by:

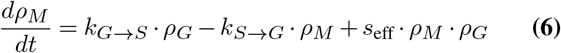

where *k*_*G*→*S*_ and *k*_*S*→*G*_ are the switching rates (Eqs. 1-2) and *s*_eff_ = *b*_*m*_ − *b*_*wt*_ is an effective selection coefficient that captures the fitness difference in a well-mixed (mean-field) approximation. At steady state (*dρ*_*M*_ */dt* = 0), and substituting *ρ*_*G*_ = 1 − *ρ*_*M*_, this yields a quadratic equation for *ρ*_*M*_. Solving for *ρ*_*M*_ gives the theoretical steady-state fraction:

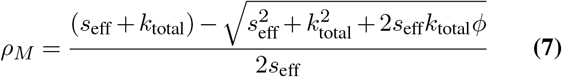

For *s*_eff_ = 0, this simplifies to *ρ*_*M*_ = (1 −*ϕ*)*/*2. In Fig. S4, we fit *s*_eff_ as a single free parameter to the agent-based simulation data for each *b*_*m*_ value.

## ACKNOWLEDGEMENTS

V.S. gratefully acknowledges the support of the ”Scholarships for Meritorious Students to Study Abroad”, offered by the Higher Education Department, Government of Madhya Pradesh, India, and the Pundit RD Sharma Memorial Graduate Award in Mathematical and Statistical Sciences. The research of H. W. was partially supported by the Natural Sciences and Engineering Research Council of Canada (NSERC) (Individual Discovery Grant RGPIN-2025-05734, Discovery Accelerator Supplement Award RGPAS-2020-00090) and the Canada Research Chairs program (Tier 1 Canada Research Chair Award). The research of both V.S. and H.W. was partially supported by the NSERC Alliance Missions Grant 577242. This research was enabled in part by support provided by Calcul Québec (calculquebec.ca), the BC DRI Group (bcdri.ca), and the Digital Research Alliance of Canada (alliancecan.ca).

## Data Availability

The custom simulation code is available on Github. The datasets generated and analyzed during the current study are available from the corresponding author upon reasonable request.

## Supplementary Information

**Fig. S1.**
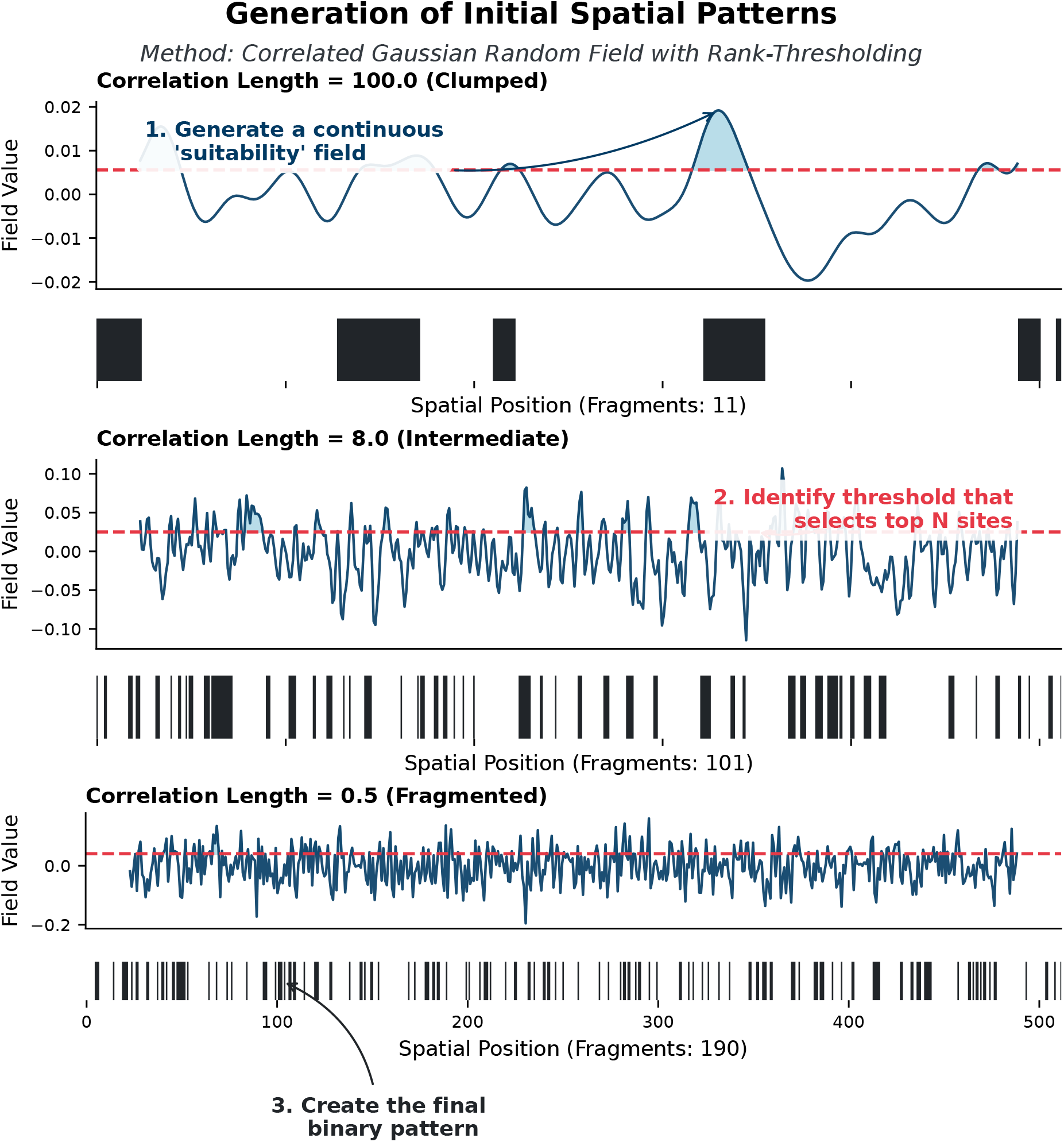
Method for Generating Spatially Correlated Initial Conditions. To control the initial spatial arrangement of specialists, we use a method based on a Correlated Gaussian Random Field (GRF). (1) A continuous 1D field of ”suitability” values is generated. The smoothness of this field is controlled by the correlation length parameter. (2) A rank-based threshold is applied to select the top *N*_*S*_ sites with the highest field values (shaded region above the red dashed line). (3) These sites are converted into a binary pattern of Specialist (black) and Generalist (white) cells. A large correlation length (top) results in a single clump, while a small correlation length (bottom) results in a highly fragmented pattern, all while keeping the total number of specialists constant.

**Fig. S2.**
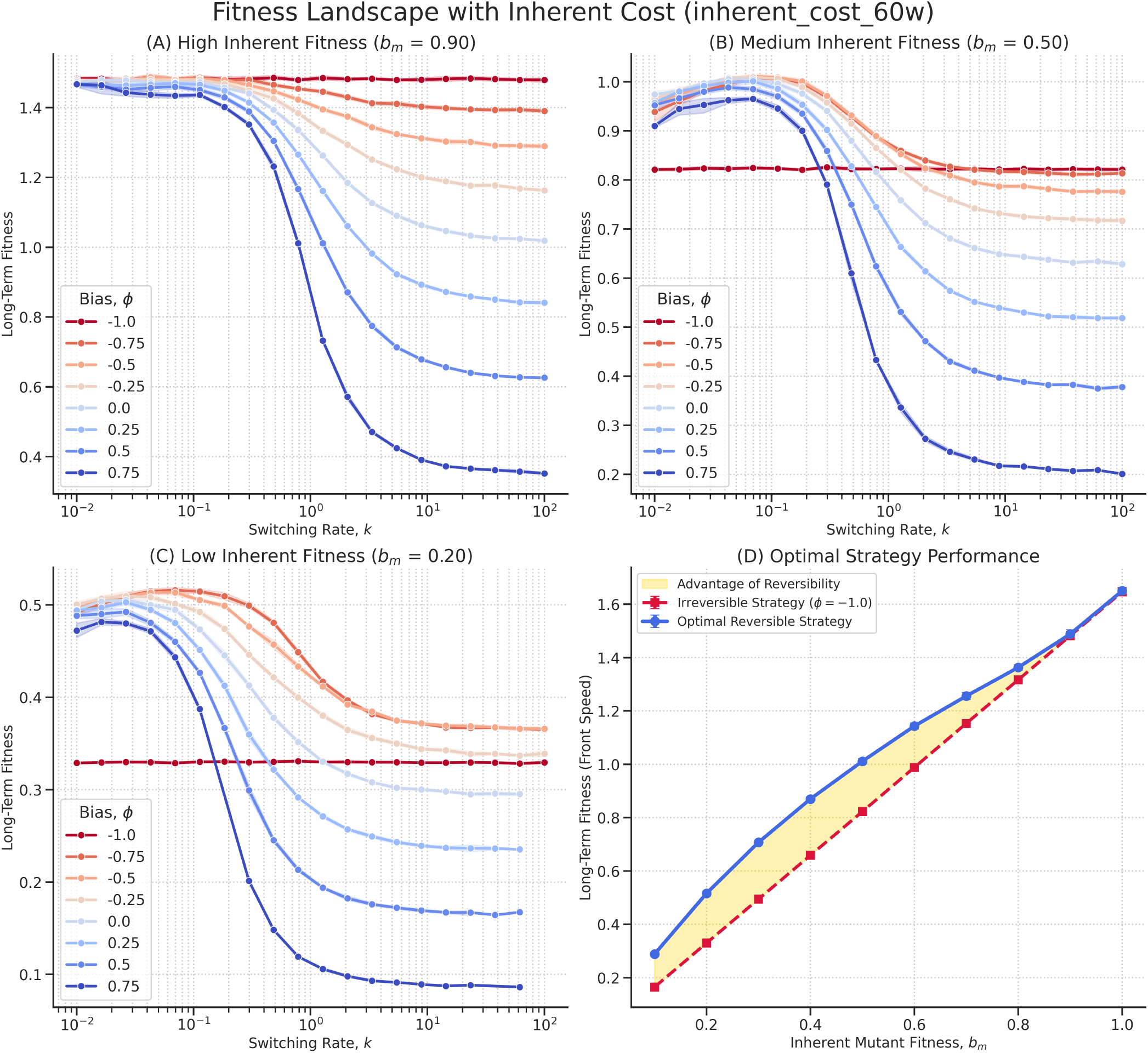
Reversibility Confers a Fitness Advantage in the ”Resilience” Trade-off. This figure replicates the analysis from Fig. 2 of the main text, but for a ”resilience” trade-off analogous to bacterial persistence, where the specialist phenotype incurs a fixed, inherent fitness cost in all environments (here, *b*_*m*_ is always less than *b*_*wt*_ = 1.0). (A-C) Fitness landscapes as a function of switching rate (*k*_total_) and bias (*ϕ*) for different magnitudes of the specialist’s fitness cost. Reversible strategies (blue-toned lines, *ϕ >* −1.0) consistently outperform the irreversible strategy (red line, *ϕ* = −1.0). (D) The performance of the optimal strategy for each condition. The yellow shaded area highlights the ”Advantage of Reversibility”—the fitness gained by allowing back-switching. Even when the specialist phenotype is never optimal, a flexible, reversible strategy remains superior, demonstrating the generality of this principle.

**Fig. S3.**
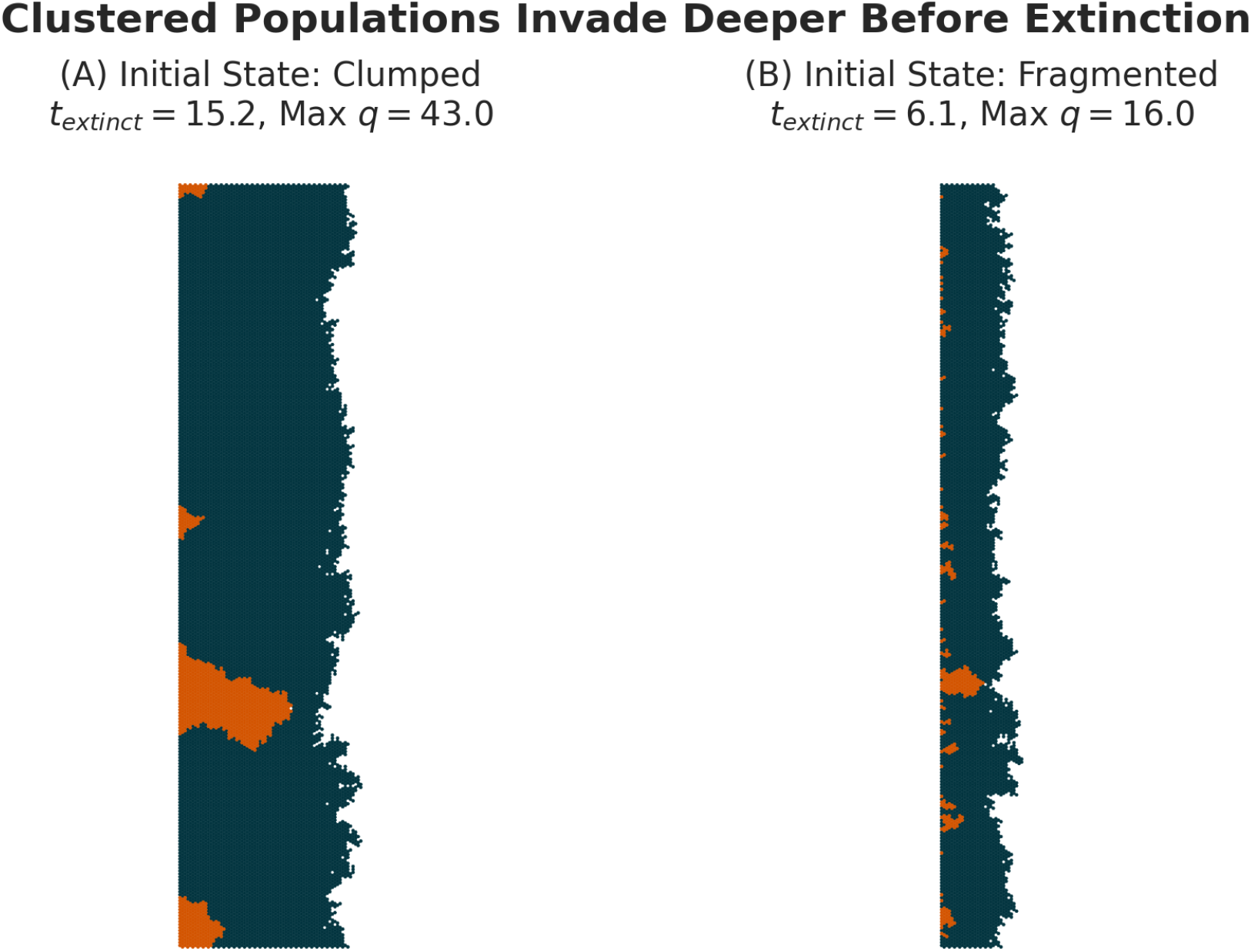
Direct Visualization of Collective Protection. This figure shows the final state of two simulations run until extinction, both starting with the same number of disadvantaged specialists (*b*_*m*_ = 0.7) but with different initial spatial arrangements. (A) When specialists start in a single large cluster, they are able to collectively shield individuals in the interior from selection, allowing the lineage to invade deep into the territory before eventually going extinct. (B) When the same number of specialists are highly fragmented, each small cluster or individual has a large perimeter-to-area ratio, exposing them to strong negative selection from the surrounding generalists. This leads to rapid, wholesale extinction with minimal spatial expansion. This provides a direct visual confirmation of the mechanism underlying the results in Fig. 5 of the main text.

**Fig. S4.**
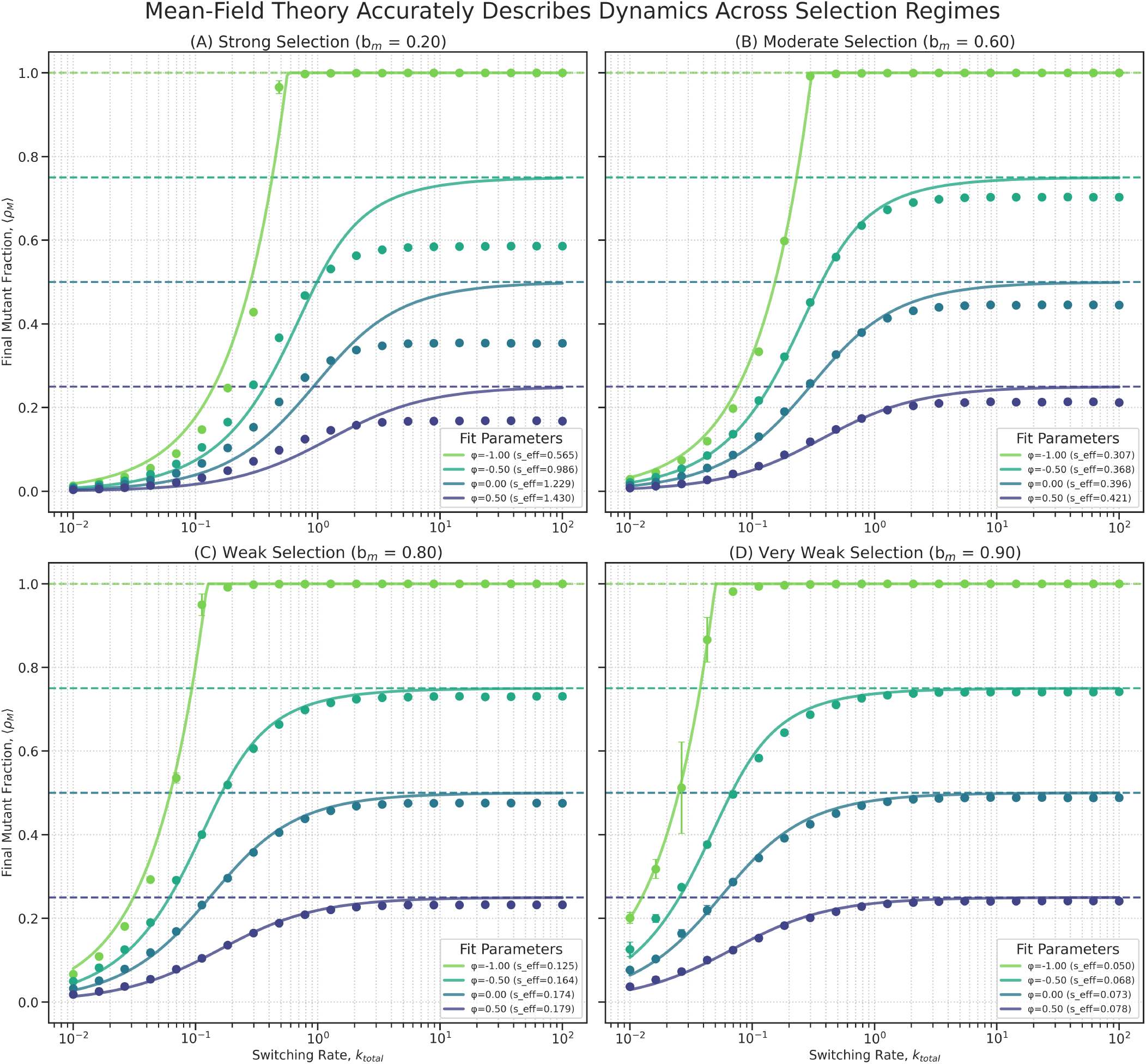
Mean-field theory accurately describes steady-state dynamics across selection regimes. This figure validates the mean-field model described in the Methods section against the full agent-based model. Each panel shows the final specialist fraction ⟨*ρ*_*M*_ ⟩ as a function of switching rate (*k*_total_) for a different selection strength (*b*_*m*_). **Points (with error bars):** Mean ± standard deviation from the full agent-based simulations. **Solid lines:** Fits of the mean-field model (Eq. 7), using *s*_eff_ as a single fitting parameter. **Dashed lines:** The neutral expectation *ρ*_*M*_ = (1 − *ϕ*)*/*2, which is the high-*k*_Gtotal_ asymptote. The model fits the simulation data well, especially for weak selection (C, D), but begins to deviate under strong selection (A), confirming that the mean-field model is a good approximation for the final steady state but not a perfect representation of the full spatial dynamics.

**Fig. S5.**
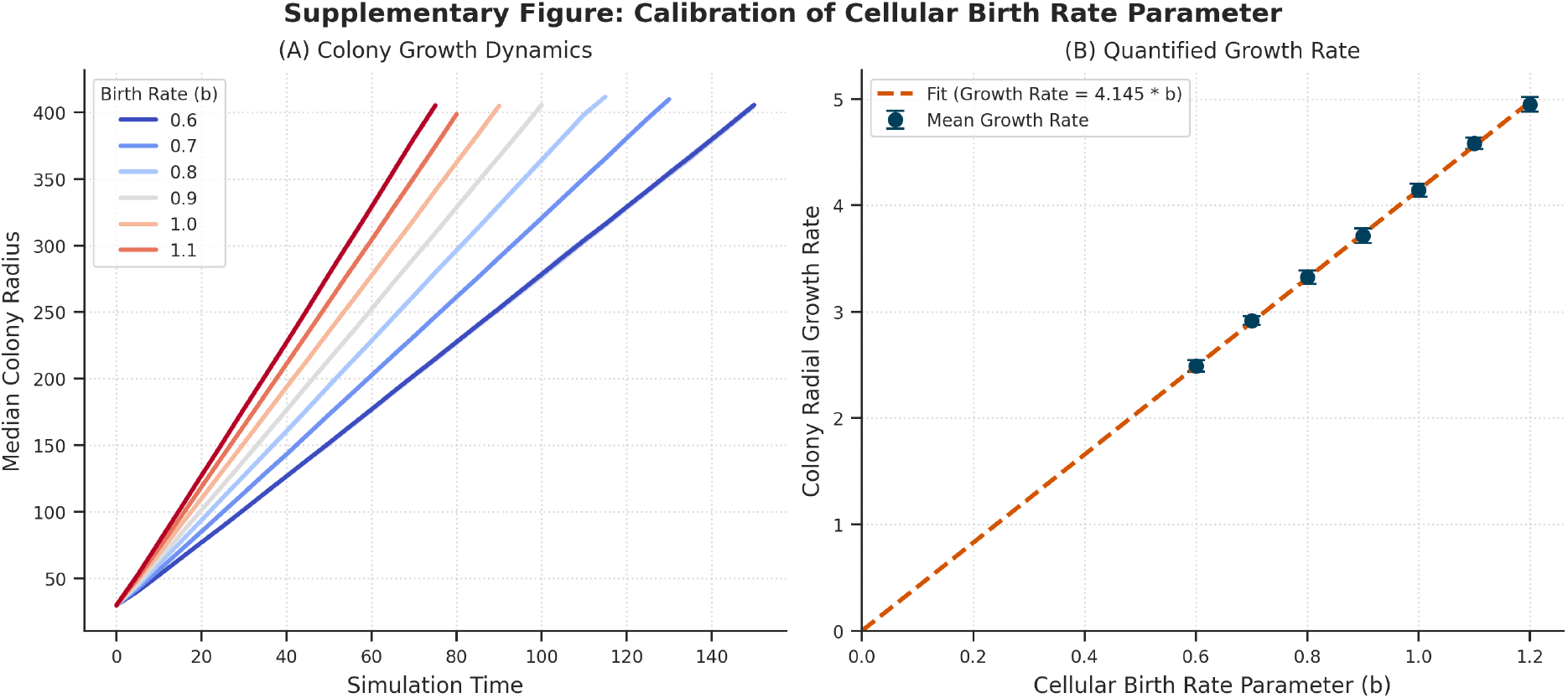
Calibration of Cellular Birth Rate Parameter. This figure demonstrates the relationship between the input cellular birth rate parameter (*b*) used in the agent-based model and the resulting macroscopic radial growth rate of the colony. **(A) Colony Growth Dynamics:** Median colony radius over simulation time for various values of *b*. The growth is approximately linear after an initial transient. **(B) Quantified Growth Rate:** The mean radial growth rate (slope extracted from panel A) plotted against the input parameter *b*. Error bars represent the standard deviation across replicates. The dashed line shows a linear fit constrained to pass through the origin (Growth Rate = constant × *b*), confirming the expected proportionality and providing the calibration constant used to relate microscopic parameters to macroscopic observables.

**Fig. S6.**
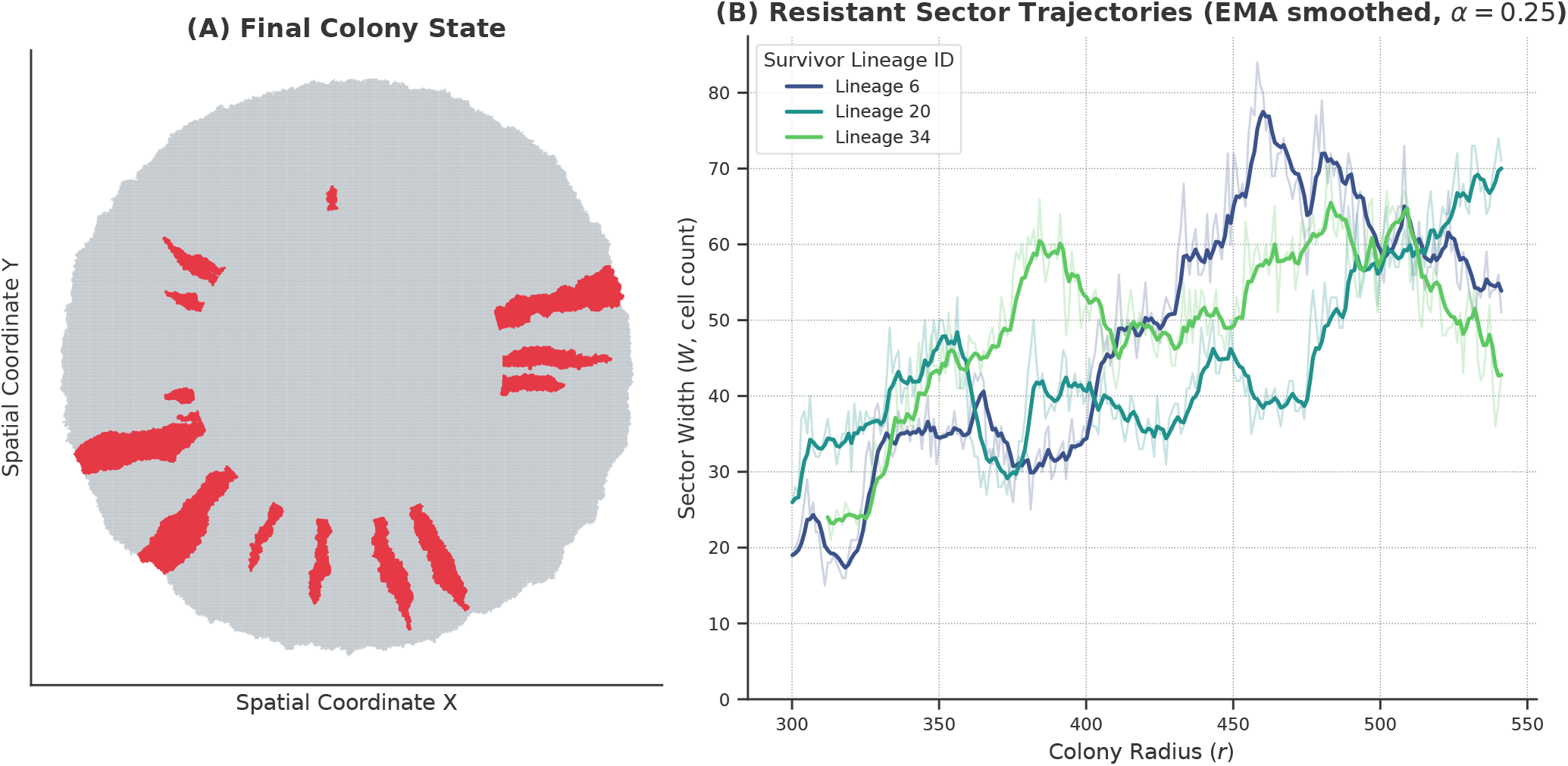
Illustration of Sector Width Dynamics in Radial Expansion. This figure shows an example output from the radial expansion simulation used for the analysis in Fig. 6 (main text), illustrating the sectoring dynamics and the online width tracking method. **(A) Full Colony View:** The final state of the simulated colony, starting from multiple small resistant sectors (red) in a wild-type background (light gray). Red cells indicate the resistant phenotype. **(B) Survivor Sector Widths:** The width (in cells) of three representative resistant sectors that survived until the end of the simulation, plotted against colony radius. Widths are measured dynamically at the front during the simulation (”Online widths”). Both raw data (thin lines) and smoothed data (thicker lines, using exponential moving average) are shown, demonstrating the significant fluctuations inherent in boundary dynamics at the rough front.

**Fig. S7.**
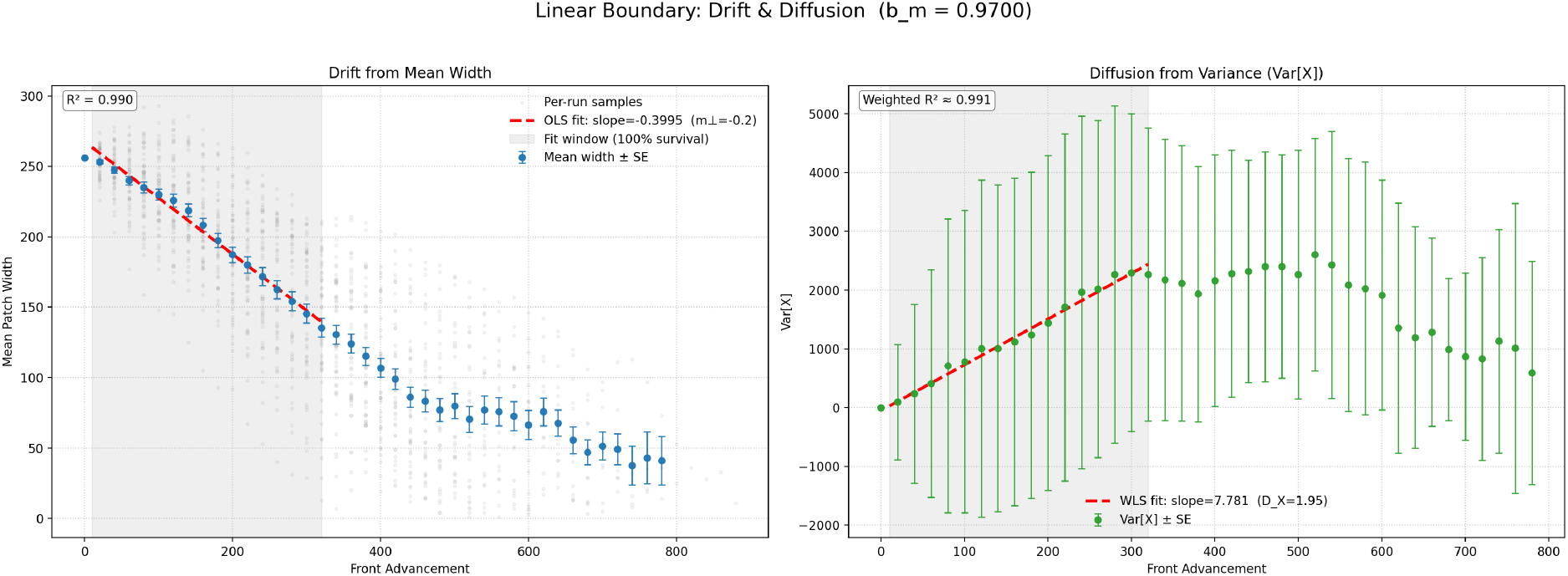
Inference of Conditioned Random Walk (RW) Model Parameters. This figure illustrates the parameter inference procedure described in the Methods and used for the linear expansion case in Fig. 6A (*b*_*m*_ ≈ 0.97, *W*_0_ = 256). **Left (Drift):** The effective drift *m*_⊥_ is inferred from the slope of an Ordinary Least Squares (OLS) linear regression on the mean patch width vs. front advancement (Eq. 4). **Right (Diffusion):** The effective diffusion *D*_*X*_ is inferred from the slope of a Weighted Least Squares (WLS) linear regression on the patch width variance vs. front advancement (Eq. 5).

